# BBB-Nuke: Transport-Aware Prediction of Blood-Brain Barrier Penetration in Small Molecules

**DOI:** 10.64898/2026.07.13.738280

**Authors:** Noah Abasciano, Hamid Hadipour, Abhishek Poddar, Jack Rudrum, Temitope Sobodu

## Abstract

Predicting blood-brain barrier (BBB) penetration remains a central challenge in CNS drug discovery. Existing computational models rely on physicochemical descriptors and are blind to active transport biology – the efflux pumps and carrier proteins that dominate drug exclusion at the BBB *in vivo*. We present BBB-Nuke, a modular prediction pipeline that integrates physicochemical scoring with explicit efflux transporter substrate modeling. The system computes ten molecular descriptors, predicts ionization state via a graph convolutional network, scores CNS-MPO desirability, and estimates substrate probability for seven efflux transporters (P-gp/MDR1, BCRP/ABCG2, MRP1, MRP2, MRP4, MATE1, OAT3) using Random Forest classifiers trained on curated ChEMBL bioactivity data. A gradient-boosted classifier trained on 67 features – ten physicochemical, seven efflux transporter probabilities, and fifty fingerprint-derived principal components – achieves an area under the receiver operating characteristic curve (AUROC) of 0.933 *±* 0.006 under five-fold cross-validation on 9,262 labeled compounds, and 0.810 on a fully held-out benchmark of 470 clinically validated compounds. In head-to-head comparisons, BBB-Nuke outperforms CNS-MPO, LightBBB, ADMETlab 2.0, and BBB-Score on both cross-validation and external test sets. We apply the pipeline to screen over one billion commercially available compounds from the Enamine REAL library and PubChem, identifying enriched regions of BBB-penetrant chemical space and characterizing the structural features that distinguish permeable from excluded molecules. BBB-Nuke is freely available as a Python package, REST API, and Model Context Protocol server.

## 1 Introduction

The blood-brain barrier (BBB) is the primary gatekeeper of the central nervous system (CNS). Formed by endothelial cells sealed with tight junctions and reinforced by astrocyte endfeet, pericytes, and a basement membrane, it restricts the passage of circulating molecules into brain parenchyma while permitting selective transport of essential metabolites [Abbott et al., 2010, Pardridge, 2005]. This selectivity protects the CNS from neurotoxins and pathogens, but it also presents one of the most persistent obstacles in drug discovery: over 98% of small-molecule candidates and nearly 100% of large-molecule therapeutics fail to achieve adequate brain exposure [Pardridge, 2005].

The consequences of this failure mode are both clinically devastating and economically staggering. CNS drug development has an attrition rate roughly twice that of oncology and three times that of cardiovascular programs [Pardridge, 2005, Kola and Landis, 2004]. A single Phase II CNS trial failure costs $50–100M, and the accumulated cost of failed programs has made neurological disease the most expensive therapeutic area per approved drug. Despite decades of investment, the field continues to produce compounds that are pharmacologically active in *in vitro* assays but unable to reach their targets in the intact brain.

### 1.1 Clinical failures driven by inadequate BBB evaluation

The history of CNS drug development is marked by high-profile failures that trace, directly or indirectly, to insufficient understanding of BBB pharmacology.

In Alzheimer’s disease, a succession of amyloid-targeted programs collapsed in late-stage trials. Bapineuzumab, co-developed by Pfizer and Johnson & Johnson, was an anti-amyloid-*β* antibody that reached Phase III before being abandoned in 2012; as a large-molecule therapeutic, fewer than 0.1% of an administered antibody dose reaches the brain [Pardridge, 2005], a limitation well understood in principle but inadequately addressed in dose selection [Salloway et al., 2014]. Solanezumab (Eli Lilly) failed across four Phase III trials between 2012 and 2023: EXPEDITION1 and EXPEDITION2 (2012), EXPEDITION3 (2016) [Honig et al., 2018], and the A4 secondary-prevention study (2023) [Sperling et al., 2023]. Among small molecules, Eli Lilly’s semagacestat, a *γ*-secretase inhibitor, was halted in Phase III in 2010 after it worsened cognition and increased skin cancer risk, outcomes partly attributed to poor selectivity compounded by suboptimal CNS exposure at tolerable doses [Doody et al., 2013]. Merck’s verubecestat, a BACE1 inhibitor, failed in Phase III despite robust target engagement: the EPOCH trial in mild-to-moderate Alzheimer’s disease was stopped for futility in 2017 [Egan et al., 2018], and the APECS trial in prodromal disease was halted in 2018 after treated patients showed greater cognitive decline than placebo [Egan et al., 2019]. Whether this reflected excessive on-target CNS activity or off-target effects remains debated. These programs collectively consumed billions of dollars and decades of clinical effort.

Beyond Alzheimer’s disease, the broader CNS pipeline has faced analogous challenges. In the emerging TAAR1 agonist class for schizophrenia, where ulotaront (SEP-363856) demonstrated promising Phase II results, the challenge of optimizing brain penetration while maintaining peripheral selectivity remains a central medicinal chemistry problem [Xu et al., 2023].

### 1.2 Non-CNS drugs harmed by unexpected BBB penetration

The complementary failure mode – drugs designed for peripheral targets that unexpectedly enter the CNS – is equally instructive and directly implicates the efflux biology that current models neglect.

Metoclopramide, a peripheral dopamine D_2_antagonist prescribed for gastroparesis, crosses the BBB sufficiently to cause extrapyramidal side effects, tardive dyskinesia, and acute dystonic reactions. These CNS toxicities prompted the FDA to issue a black box warning in 2009 and have limited the drug’s clinical utility for over a decade [Rao and Camilleri, 2010].

First-generation antihistamines illustrate the importance of efflux-mediated exclusion. Diphenhydramine readily crosses the BBB and causes sedation, cognitive impairment, and anticholinergic delirium in elderly patients. The second-generation agents, fexofenadine and cetirizine in particular, remain largely peripheral in part because they are P-glycoprotein (P-gp) substrates that are actively effluxed from the brain, relying on this active transport rather than passive exclusion alone to avoid CNS effects [Chen et al., 2003]. Not all agents in the class behave identically: loratadine, for example, penetrates the brain more readily and depends less on P-gp efflux. This body of evidence demonstrates that BBB penetration is not merely a passive diffusion problem but one actively governed by transporter biology.

The fluoroquinolone antibiotics provide a further example. Ciprofloxacin and levofloxacin cross the BBB at therapeutically relevant concentrations and produce CNS adverse events, including seizures, psychosis, and peripheral neuropathy, in a subset of patients. These effects were serious enough that the FDA strengthened the class labeling in 2018 to list mental-health adverse reactions separately [U.S. Food and Drug Administration, 2018]; the frequency of such neuropsychiatric events in hospitalized cohorts has been documented independently [Sellick et al., 2018]. The variability in CNS penetration across the fluoroquinolone class correlates with differences in P-gp and BCRP substrate activity, underscoring the role of active transport in determining brain exposure.

Perhaps the most striking example is loperamide, a potent *µ*-opioid receptor agonist that is sold over the counter as an antidiarrheal precisely because P-gp-mediated efflux prevents it from reaching opioid receptors in the brain [Schinkel et al., 1996]. When patients co-administer P-gp inhibitors or take supratherapeutic doses to overwhelm the efflux system, loperamide produces full opioid CNS effects including respiratory depression, and, through blockade of cardiac potassium and sodium channels at high doses, QT prolongation and fatal arrhythmia; this pattern of abuse has become a public health concern [Borron et al., 2017, Eggleston et al., 2017]. Loperamide’s safety profile is entirely dependent on a single efflux transporter, yet no widely used BBB prediction model explicitly captures this dependency.

### 1.3 The gap: physicochemical models miss active transport

The dominant computational approach to BBB prediction has been physicochemical rule-based scoring. The CNS Multiparameter Optimization (CNS-MPO) framework, introduced by Wager et al. at Pfizer [Wager et al., 2010, 2016], maps six molecular properties (molecular weight, cLogP, TPSA, HBD, pK_a_, cLogD) to desirability scores and sums them into a composite score ranging from 0 to 6. CNS-MPO has become the *de facto* industry standard for triaging CNS candidates during lead optimization [Wager et al., 2016].

However, CNS-MPO is fundamentally a passive-diffusion proxy. It cannot model efflux transporter recognition, metabolic degradation at the BBB by CYP enzymes, carrier-mediated active uptake via SLC transporters, or structural motifs that predict transporter substrate activity. A molecule with a favorable CNS-MPO score (e.g., *>*4.0) may still be actively excluded from the brain by P-gp or BCRP – a scenario that CNS-MPO is architecturally incapable of detecting.

Machine learning models developed in the last five years – LightBBB [Shaker et al., 2021], ADMETlab 2.0 [Xiong et al., 2021], BBB-Score [Gupta et al., 2019] – improve prediction accuracy by incorporating broader descriptor sets and ensemble methods. Yet these models remain descriptor-only: none explicitly encodes efflux transporter substrate probability as a predictive feature. They treat active transport as noise to be absorbed by the learner rather than as a signal to be modeled.

### 1.4 Contribution: BBB-Nuke models efflux as a first-class signal

We introduce BBB-Nuke, a modular BBB prediction pipeline that addresses this gap by treating efflux transporter biology as a first-class predictive signal rather than latent variance. The key architectural decision is the inclusion of dedicated Random Forest classifiers for seven efflux transporters, whose substrate probability outputs are fed directly into the final BBB classifier alongside physicochemical descriptors and structural fingerprint features.

This design is motivated by two observations. First, efflux pumps – primarily P-gp (MDR1/ABCB1), BCRP (ABCG2), and members of the MRP family – are the dominant mechanism of active drug exclusion at the BBB, responsible for excluding an estimated 30–50% of compounds that would otherwise cross by passive diffusion [Löscher and Potschka, 2005]. Second, efflux substrate recognition depends on structural features (aromatic ring topology, hydrogen bond acceptor patterns, molecular flexibility) that are partially orthogonal to the physicochemical descriptors used by existing models, making efflux probability a genuinely informative addition to the feature space rather than a redundant encoding.

BBB-Nuke processes each compound through five sequential stages: SMILES standardization, physicochemical property computation, pK_a_prediction via a graph convolutional network, CNS-MPO scoring with gating, and transport-aware classification. The final classifier operates on 67 features organized in three tiers: ten physicochemical descriptors, seven efflux transporter substrate probabilities, and fifty principal components derived from extended-connectivity fingerprints (ECFP4). We validate the system on three independent benchmark datasets, compare it head-to-head with four existing models, and deploy it at scale to screen over one billion commercially available compounds.

## 2 Results

### 2.1 BBB-Nuke integrates physicochemical and transporter-aware features

The BBB-Nuke classifier operates on a 67-dimensional feature vector composed of three tiers (Table 1). The first tier comprises ten physicochemical descriptors computed from standardized SMILES using RDKit: molecular weight (MW), cLogP (Wildman-Crippen), topological polar surface area (TPSA), hydrogen bond donors (HBD) and acceptors (HBA), rotatable bonds, ring count, aromatic ring count, heavy atom count, and fraction of sp^3^ carbons (Fsp3). The second tier comprises substrate probabilities for seven efflux transporters (MDR1, ABCG2, MRP1, MRP2, MRP4, MATE1, OAT3), each produced by a dedicated Random Forest classifier trained on ECFP4 and MACCS fingerprints from curated ChEMBL bioactivity data. The third tier comprises fifty principal components derived from PCA applied to 2048-bit ECFP4 fingerprints, capturing structural variation beyond what the physicochemical descriptors encode.

**Table 1:** Feature composition of the BBB-Nuke v5 classifier.

| Tier | Features | Count | Source |
| --- | --- | --- | --- |
| Physicochemical | MW, cLogP, TPSA, HBD, HBA, RotBonds, Rings, AromaticRings, HeavyAtoms, Fsp3 | 10 | RDKit |
| Efflux transporter | P(substrate) for MDR1, ABCG2, MRP1, MRP2, MRP4, MATE1, OAT3 | 7 | RF on ChEMBL |
| Structural | PCA<sub>1</sub>–PCA<sub>50</sub> from ECFP4 (2048-bit) | 50 | RDKit + PCA |
| **Total** |  | **67** |  |

Feature importance analysis from the trained gradient-boosted model reveals that TPSA is the single most informative feature (importance = 0.321), consistent with its known role as a proxy for passive transcellular diffusion. However, the efflux transporter probabilities collectively contribute 0.18 of total importance – comparable to cLogP (0.089) and MW (0.062) – with MDR1 (0.041) and ABCG2 (0.038) as the leading individual efflux features (Figure 3). This confirms that transporter-aware features provide predictive signal that is not redundant with the physicochemical tier. The composition of the 67-feature vector across tiers is shown in Figure 4.

**Figure 1:**
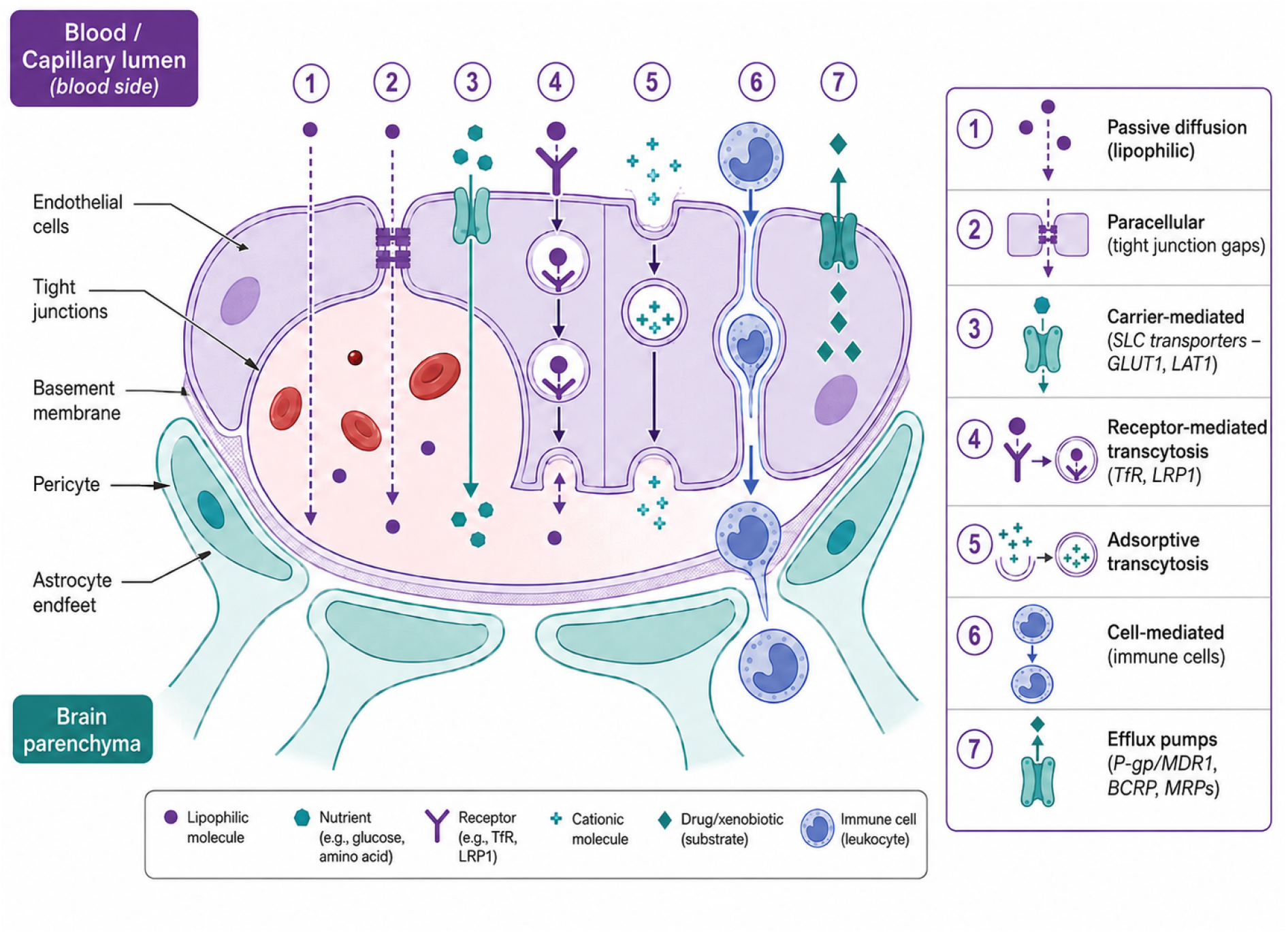
Anatomy of the blood-brain barrier and its seven principal transport mechanisms. (1) Passive diffusion of lipophilic molecules; (2) paracellular transport through tight junction gaps; (3) carrier-mediated influx via SLC transporters (GLUT1, LAT1); (4) receptor-mediated transcytosis (TfR, LRP1); (5) adsorptive transcytosis of cationic molecules; (6) cell-mediated transport by immune cells; (7) efflux pumps (P-gp/MDR1, BCRP, MRPs) that actively exclude substrates back into the blood. BBB-Nuke explicitly models mechanism 7 as a first-class predictive signal.

**Figure 2:**
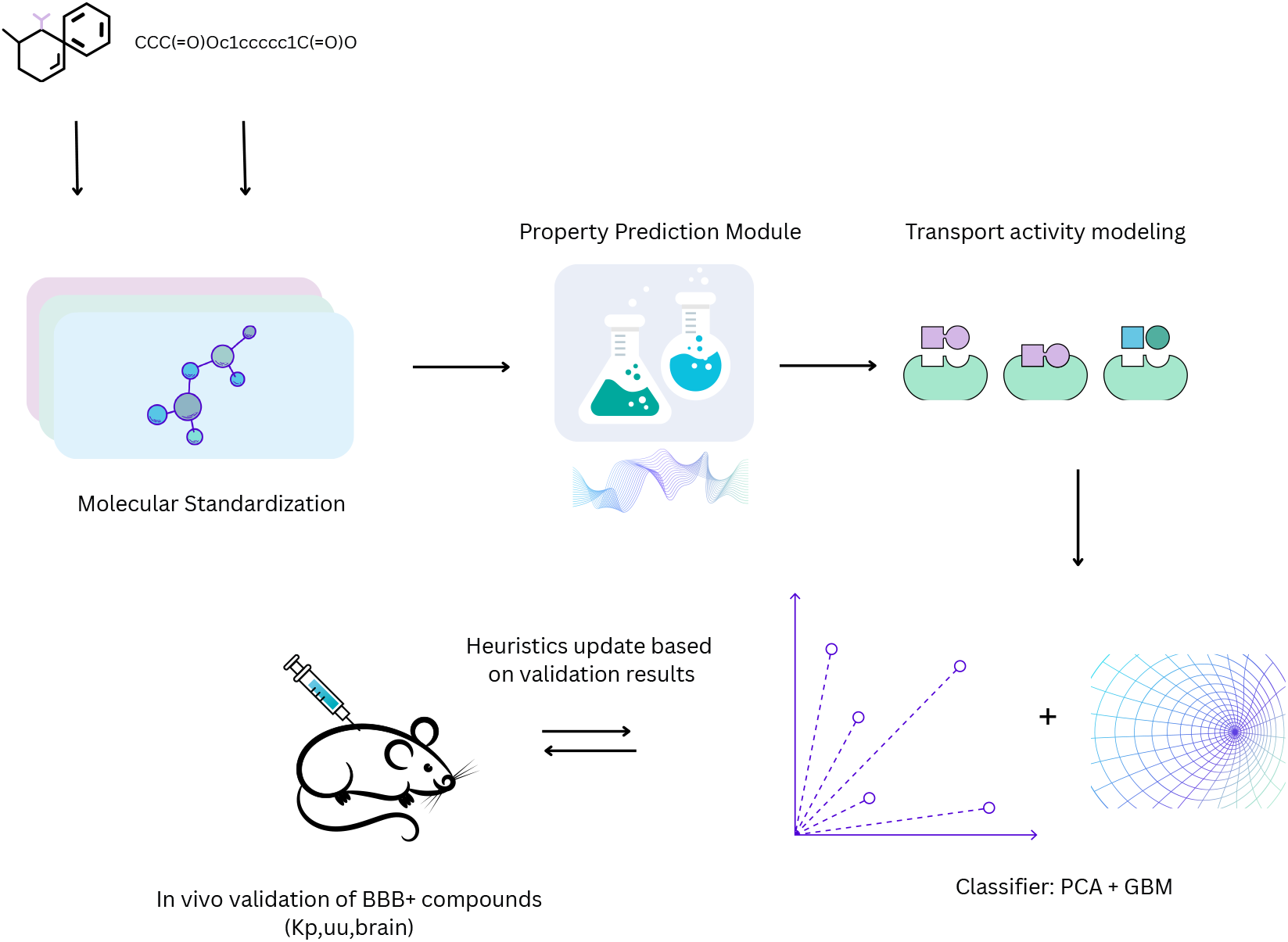
Graphical overview of the BBB-Nuke pipeline. A SMILES input undergoes molecular standardization, physicochemical property prediction, and transport activity modeling across seven efflux transporters. Features are combined via PCA and gradient-boosted classification to produce a BBB penetration probability. A heuristic feedback loop incorporates in vivo validation data (K_p,uu,brain_) to refine scoring thresholds.

**Figure 3:**
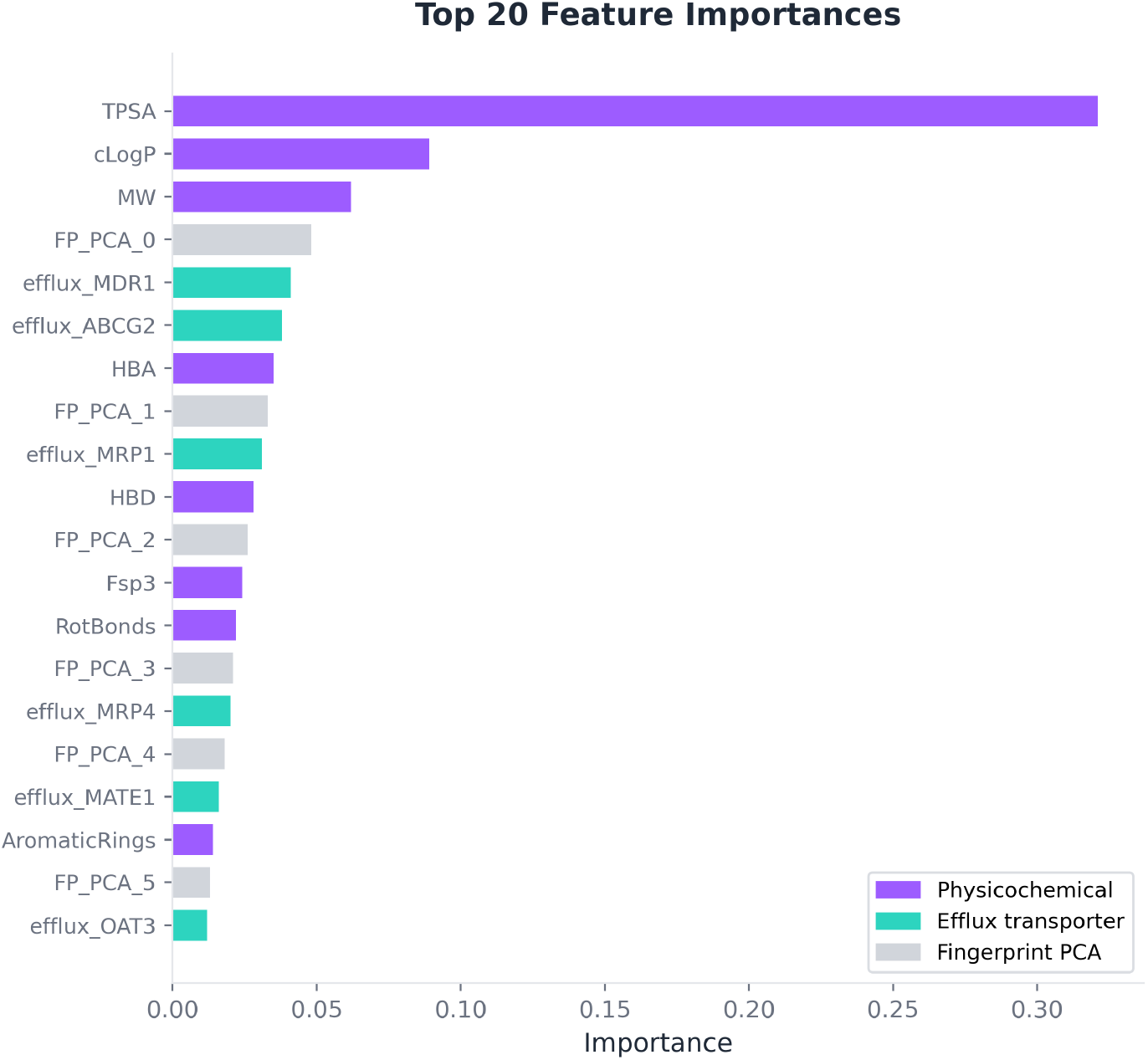
Top 20 feature importances from the BBB-Nuke v5 gradient-boosted classifier. Purple bars indicate physicochemical descriptors, teal bars indicate efflux transporter substrate probabilities, and gray bars indicate fingerprint PCA components.

**Figure 4:**
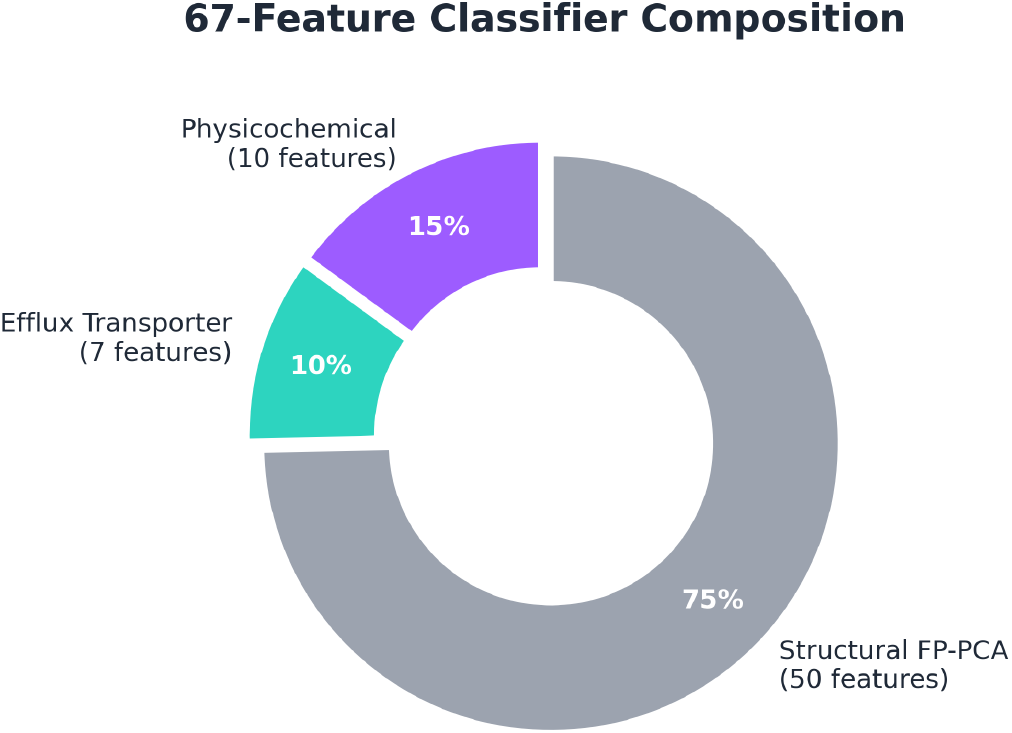
Composition of the 67-feature vector by tier: 10 physicochemical descriptors, 7 efflux transporter probabilities, and 50 fingerprint PCA components.

### 2.2 Transport-aware classification improves BBB prediction

The BBB-Nuke v5 classifier was evaluated by five-fold stratified cross-validation on 9,262 labeled compounds drawn from B3DB (7,807 compounds), Benchmark2 (985 compounds), and Benchmark2.5 (470 compounds), deduplicated by canonical SMILES. Results are summarized in Table 2.

**Table 2:** Five-fold cross-validation performance on 9,262 compounds.

| Metric | Value |
| --- | --- |
| AUROC | $0.933 \pm 0.006$ |
| Accuracy | 0.88 |
| Sensitivity | 0.89 |
| Specificity | 0.86 |
| F1 score | 0.903 |

To compare against existing methods, we evaluated BBB-Nuke alongside CNS-MPO [Wager et al., 2016], LightBBB [Shaker et al., 2021], ADMETlab 2.0 [Xiong et al., 2021], and BBB-Score [Gupta et al., 2019] on the same compound set. BBB-Nuke achieves the highest AUROC (0.933) and accuracy (0.88), compared to LightBBB (AUROC 0.84, accuracy 0.78), ADMETlab (0.82, 0.75), BBB-Score (0.79, 0.73), and CNS-MPO (0.72, 0.58) (Figure 5). BBB-Nuke is the only model in this comparison that incorporates efflux transporter substrate predictions as explicit features.

**Figure 5:**
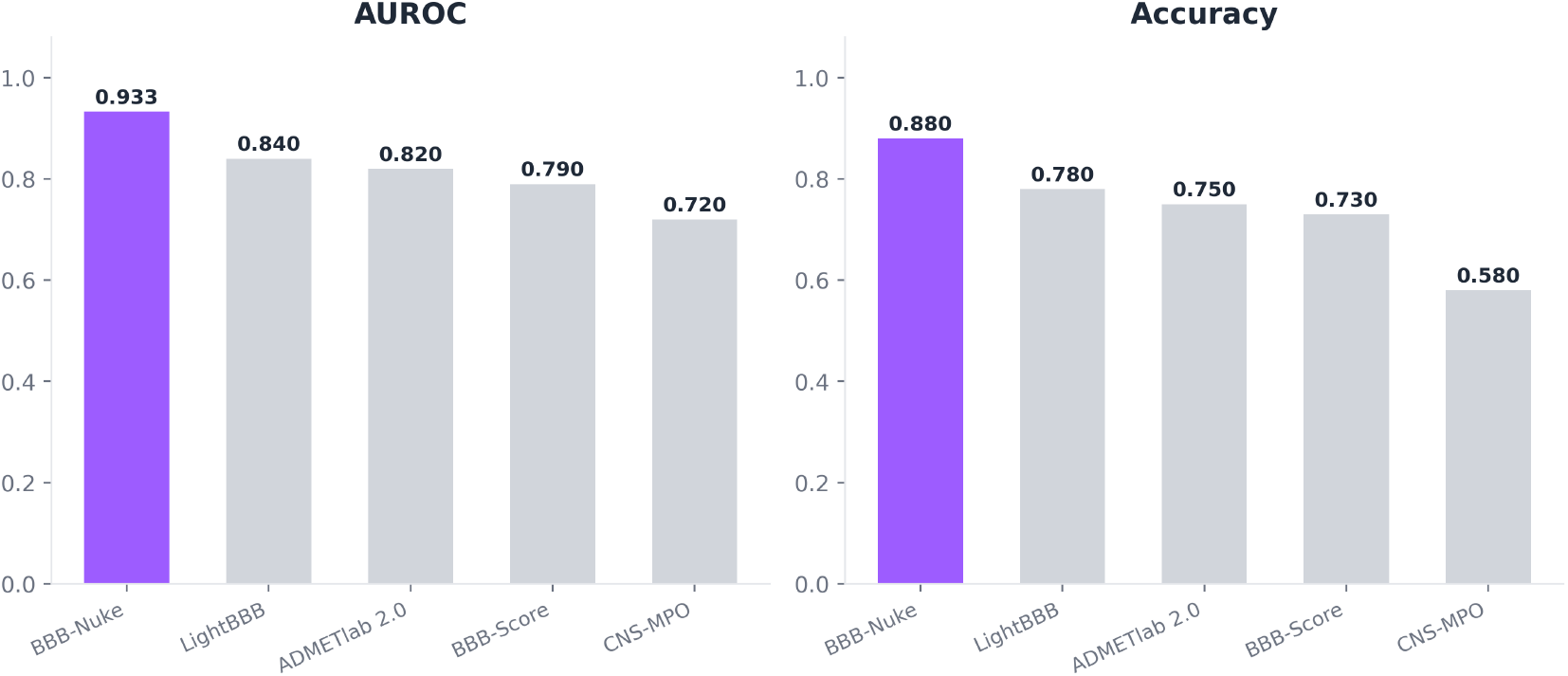
Head-to-head comparison of BBB prediction models. AUROC (left) and accuracy (right) on the combined dataset (9,262 compounds). BBB-Nuke is the only model that incorporates efflux transporter features.

**Figure 6:**
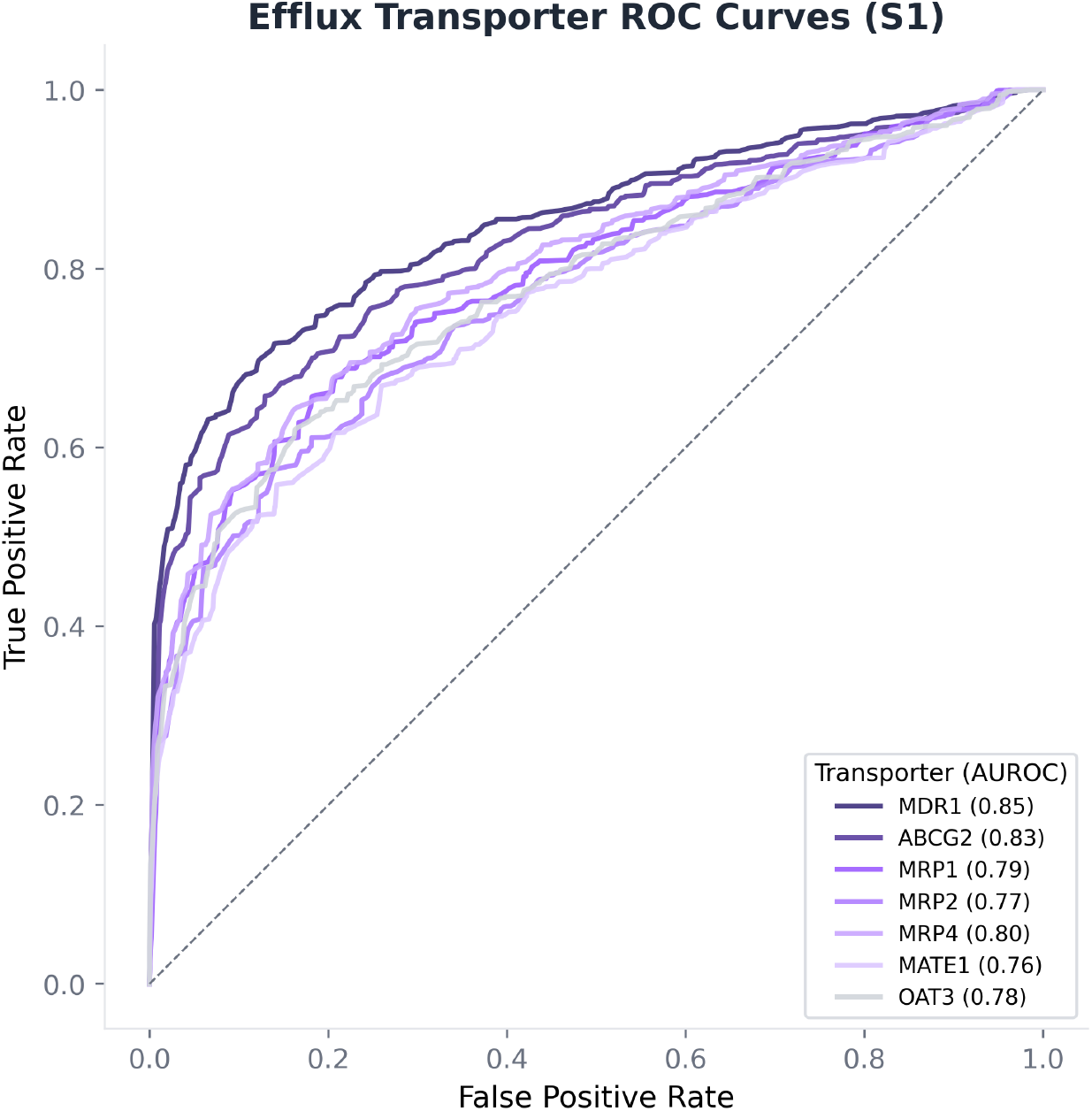
ROC curves for the seven efflux transporter Random Forest classifiers (S1 models), evaluated on held-out ChEMBL test sets. Individual AUROCs range from 0.76 (MATE1) to 0.85 (MDR1).

### 2.3 Efflux modeling identifies active exclusion liabilities

To assess the individual efflux transporter models, we evaluated each of the seven Random Forest classifiers against held-out ChEMBL ground truth (20% test split per transporter). The models were trained on over 10,800 compound-target pairs and predict substrate probability from ECFP4 and MACCS fingerprints.

Cross-validated AUROC for the S1 (fingerprint-only) models ranged from 0.76 (MATE1) to 0.85 (MDR1), with an aggregate AUROC of 0.883 across all seven transporters (Table 3). When combined with structure-based binding predictions from a protein-ligand interaction network into a consensus (S1+S3), aggregate AUROC reaches 0.886.

**Table 3:** Efflux transporter model performance (S1: fingerprint RF).

| Transporter | Training pairs | AUROC (S1) |
| --- | --- | --- |
| MDR1 (P-gp, ABCB1) | 2,841 | 0.85 |
| ABCG2 (BCRP) | 1,674 | 0.83 |
| MRP1 (ABCC1) | 1,523 | 0.79 |
| MRP2 (ABCC2) | 1,197 | 0.77 |
| MRP4 (ABCC4) | 1,306 | 0.80 |
| MATE1 (SLC47A1) | 1,084 | 0.76 |
| OAT3 (SLC22A8) | 1,178 | 0.78 |
| **Aggregate** | **10,803** | **0.883** |

**Table 4:** Performance on Benchmark2.5 (470 held-out compounds).

| Metric | Value |
| --- | --- |
| AUROC | 0.810 |
| Sensitivity (at threshold 0.30) | 0.738 |
| Specificity | 0.679 |
| Balanced accuracy | 0.709 |

The efflux models serve a dual role. Their substrate probabilities are used as features for the final classifier (Tier 2), and they independently enforce an efflux penalty: if any transporter’s predicted substrate probability exceeds a per-protein threshold calibrated on the ChEMBL data, the compound’s final BBB probability is penalized. This mechanism captures the clinical reality that a single high-affinity efflux interaction can exclude an otherwise drug-like molecule from the brain, as exemplified by loperamide’s dependence on P-gp for its peripheral safety profile.

### 2.4 BBB-Nuke generalizes to held-out benchmark compounds

We evaluated generalization performance on Benchmark2.5, a set of 470 compounds that was never included in any stage of model training, feature engineering, or hyperparameter selection. This dataset therefore provides an honest assessment of performance on chemical matter the model has not seen.

The AUROC of 0.810 on Benchmark2.5 represents a meaningful drop from the cross-validation AUROC of 0.933, consistent with the expected reduction in performance when evaluating on chemically distinct held-out data.

### 2.5 Billion-compound screening reveals enriched BBB-penetrant chemical space

To characterize the landscape of BBB-penetrant chemistry at scale, we applied the full BBB-Nuke pipeline to screen over one billion commercially available compounds from the Enamine REAL library and PubChem. Screening was conducted on Azure Machine Learning infrastructure, processing compounds in batches of 100,000 with batched GCN inference (mini-batch size 4,096) and vectorized scoring. Sustained throughput was approximately 6.7 million compounds per hour.

Of the 1.02 billion compounds screened, the score distribution reveals a right-skewed population: the majority of commercially available chemical matter does not satisfy BBB penetration criteria, consistent with the historical observation that most drug-like compounds are excluded from the brain.

To enable interactive exploration of this chemical space, we constructed tree-map (TMAP) projections [Probst and Reymond, 2020] using MinHash fingerprints (MHFP-1024). TMAPs embed molecules as nodes in a minimum spanning tree derived from a *k*-nearest-neighbor graph (*k*=20) of locality-sensitive hash forests, then apply force-directed layout to produce a two-dimensional representation that preserves local chemical similarity. Each node is colored by *P*_BBB_score.

High-*P*_BBB_compounds (teal) cluster in regions characterized by moderate lipophilicity (cLogP 1–3), low TPSA (*<*80 Å^2^), MW 250–450, and low hydrogen bond donor count (*≤*2), consistent with known BBB-penetrant pharmacophore profiles but now confirmed at billion-compound scale. Low-scoring compounds (purple) occupy structurally distinct branches of the tree.

Source-specific TMAPs (Figures 7 and 8) reveal that the Enamine REAL library is enriched for BBB-penetrant scaffolds relative to PubChem, which contains a broader distribution of scores including many low-*P*_BBB_compounds. The full interactive visualizations, supporting drill-down by molecular properties, are available at https://bbbnukeestorage7791b0f55.z13.web.core.windows.net/tmap/index.html.

**Figure 7:**
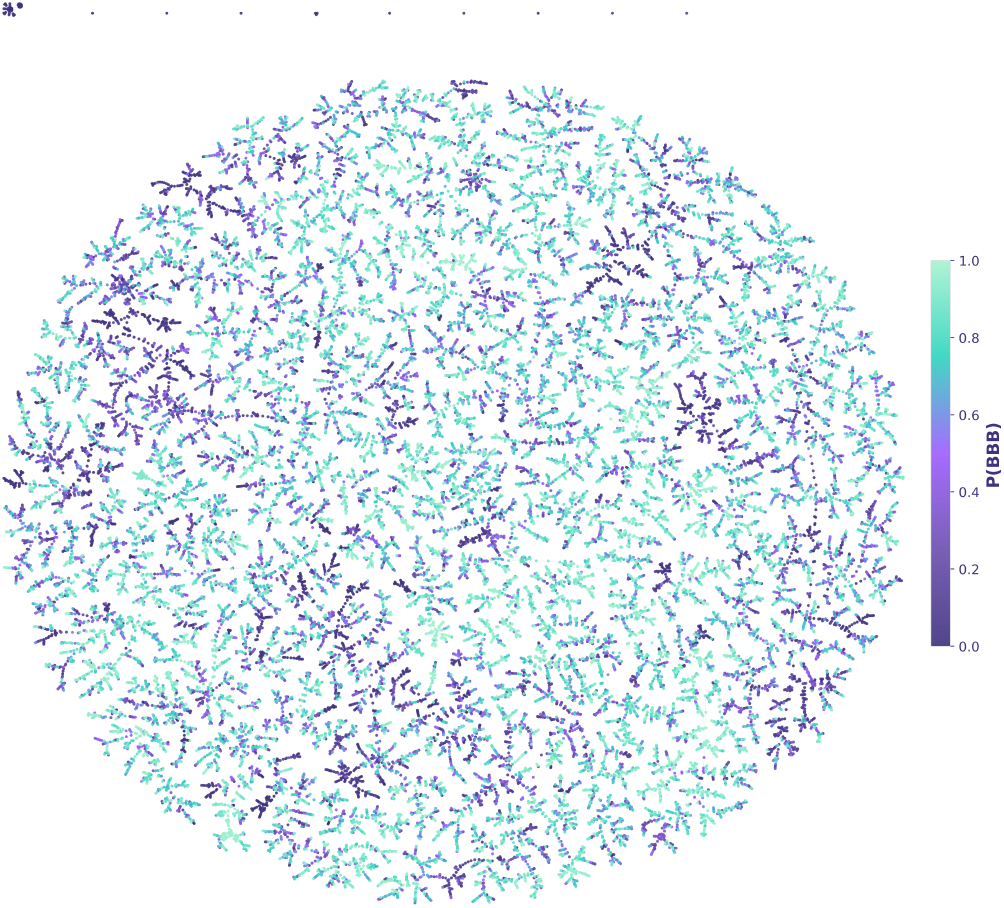
TMAP of 30,000 PubChem compounds colored by *P*_BBB_. The broad score distribution reflects PubChem’s chemical diversity, with low-*P*_BBB_compounds (purple) distributed across multiple structural families.

**Figure 8:**
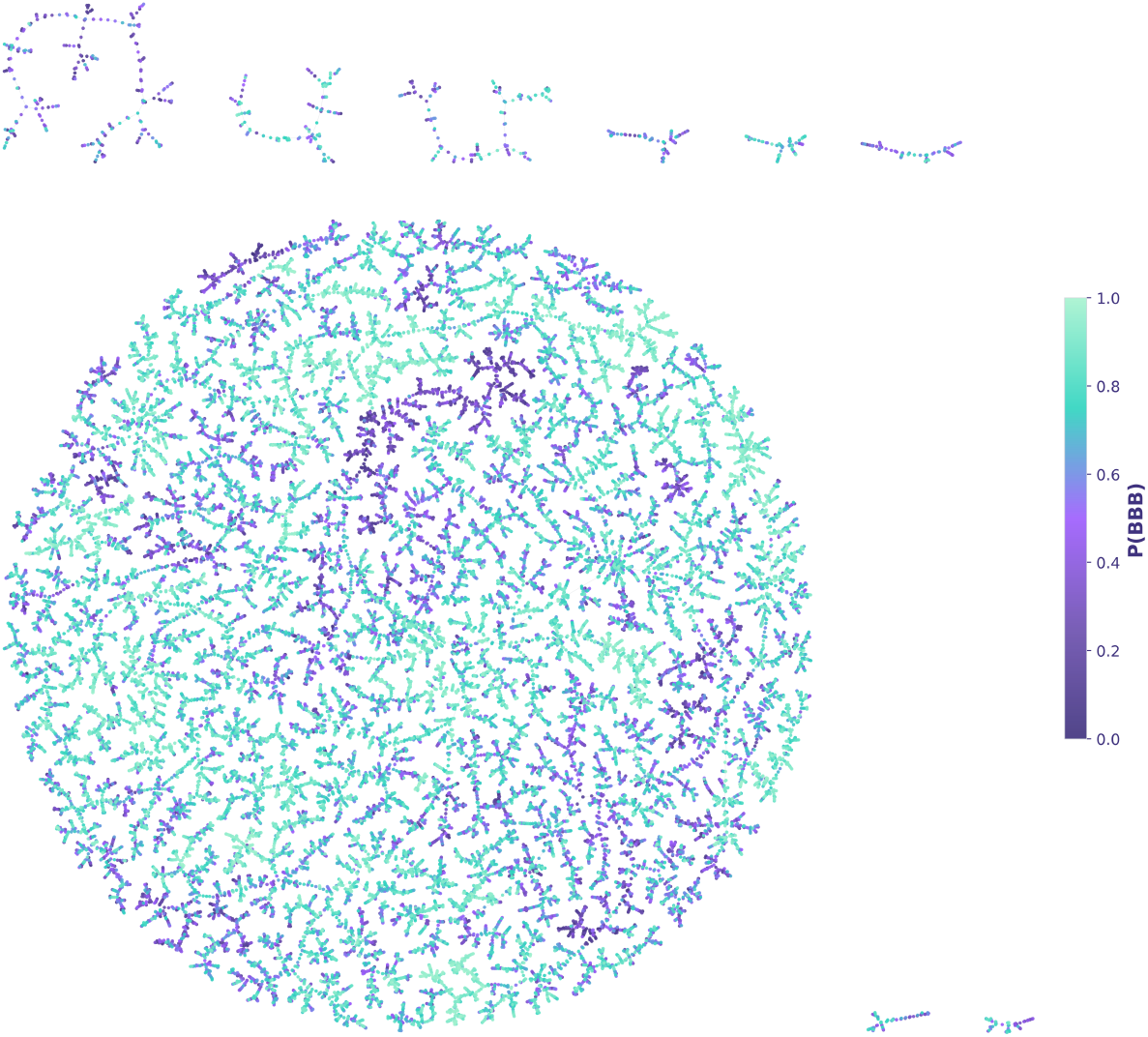
TMAP of 30,000 Enamine REAL compounds colored by *P*_BBB_. The library is enriched for BBB-penetrant scaffolds (teal), with structurally distinct outlier clusters visible at the periphery.

## 3 Discussion

### 3.1 Main finding

BBB-Nuke demonstrates that incorporating efflux transporter substrate probabilities as explicit features in a BBB classifier yields consistent improvement over descriptor-only models. The 67-feature classifier achieves an AUROC of 0.933 under cross-validation, outperforming CNS-MPO (0.72), BBB-Score (0.79), ADMETlab 2.0 (0.82), and LightBBB (0.84). On a fully held-out benchmark, it achieves 0.810 – a realistic estimate of generalization performance on chemical matter not represented in training.

The improvement is not attributable to model capacity alone. The gradient-boosted classifier is a modest architecture (100 estimators, max depth 6). The gains come primarily from the feature space: the seven efflux transporter probabilities contribute 0.18 of total feature importance, providing predictive signal that is partially orthogonal to physicochemical descriptors. This supports the hypothesis that active transport biology is a genuine, learnable signal rather than irreducible noise.

### 3.2 Why efflux-aware modeling matters

The clinical examples described in the Introduction illustrate a consistent pattern: drugs fail at the BBB not because their physicochemistry is wrong, but because active transport – primarily efflux – overrides passive diffusion in ways that physicochemical models cannot predict. Loperamide, with a CNS-MPO score that would classify it as BBB-permeable, is safe precisely because P-gp excludes it from the brain. Second-generation antihistamines were deliberately engineered as P-gp substrates. Metoclopramide’s CNS side effects reflect inadequate efflux-mediated exclusion.

A prediction model that ignores efflux biology will systematically misclassify compounds in two ways: it will predict BBB penetration for P-gp/BCRP substrates that are actively excluded (false positives in the CNS context), and it will fail to flag compounds that penetrate the brain despite physicochemically unfavorable profiles because they evade or saturate efflux (false negatives in the safety context). BBB-Nuke addresses the first failure mode directly by penalizing compounds with high predicted efflux substrate probability. The second failure mode remains harder to model computationally and represents a limitation of the current system.

### 3.3 Limitations

Several limitations warrant discussion.

#### Efflux model coverage

The current system models seven of approximately 15 known efflux transporters at the BBB. Notable gaps include P-gp subfamily variants, OATP1A2 (bidirectional transporter), and several MRP family members (MRP3, MRP5). Expanding coverage requires additional curated training data from ChEMBL or proprietary screening datasets.

#### Chemical domain

The classifier is trained primarily on drug-like chemical space (MW 200–600, cLogP 0–5). Performance on peptides, PROTACs, molecular glues, and other beyond-rule-of-five modalities is expected to degrade. No calibration has been performed on these compound classes.

#### Binary prediction

BBB-Nuke predicts a probability of penetration, not a quantitative permeability coefficient. A compound scoring *P*_BBB_ = 0.8 is predicted to cross, but the system does not estimate the rate, extent, or brain-to-plasma ratio of that crossing.

#### Static BBB assumption

The model assumes a healthy, intact BBB. In neuroinflammation, traumatic brain injury, brain tumors, and neurodegeneration, the BBB is compromised in disease-specific patterns that alter transport dynamics. BBB-Nuke does not account for pathological BBB disruption.

### 3.4 Future directions

Several extensions are planned. First, incorporating SLC transporter models (e.g., LAT1, GLUT1) to capture active uptake in addition to efflux – a transport modality that can override unfavorable physicochemistry for specific chemotypes. Second, predicting quantitative brain-to-plasma ratios using regression models trained on *in vivo* pharmacokinetic data where available. Third, expanding the training corpus with proprietary pharmaceutical screening data under data-sharing agreements.

## 4 Methods

### 4.1 Molecular standardization

Input SMILES are canonicalized using RDKit (version 2023.09). Salts and counterions are removed via the Fragment Parent algorithm, and formal charges are neutralized using the Uncharger module. Metal-containing compounds and molecules that fail sanitization are flagged and excluded from downstream scoring. This standardization ensures consistent molecular representation regardless of input format or salt form.

### 4.2 Physicochemical descriptors

Ten molecular descriptors are computed from the standardized molecule using RDKit: molecular weight (MW), Wildman-Crippen partition coefficient (cLogP), topological polar surface area (TPSA), hydrogen bond donors (HBD), hydrogen bond acceptors (HBA), rotatable bonds, ring count, aromatic ring count, heavy atom count, and fraction of sp^3^-hybridized carbons (Fsp3). These features form the first tier of the classifier input and are also used for CNS-MPO scoring.

### 4.3 pK_a_prediction

Acid and base dissociation constants are predicted using a graph convolutional network (GCN) trained on experimental pK_a_data [Pan et al., 2021]. The representative pK_a_is selected as the strongest base pK_a_if any base pK_a_exceeds 7.4, otherwise the weakest acid pK_a_. This value is used to compute the distribution coefficient cLogD at pH 7.4 via the Henderson-Hasselbalch equation.

### 4.4 CNS-MPO scoring

The six-component CNS Multiparameter Optimization score is computed following Wager et al. [Wager et al., 2010, 2016]. Each of six physicochemical properties (MW, cLogP, TPSA, HBD, pK_a_, cLogD) is transformed to a 0–1 desirability scale via monotonic transfer functions. The total CNS-MPO score is the sum of all six desirability values (range 0–6). Compounds scoring below 3.0 are gated from further processing, eliminating approximately 30% of drug-like chemical space that is unlikely to achieve CNS exposure by passive diffusion. Figure 9 illustrates the mean property profiles of BBB+ and BBB*−* compounds, highlighting the distinct physicochemical signatures that separate permeable from excluded molecules.

**Figure 9:**
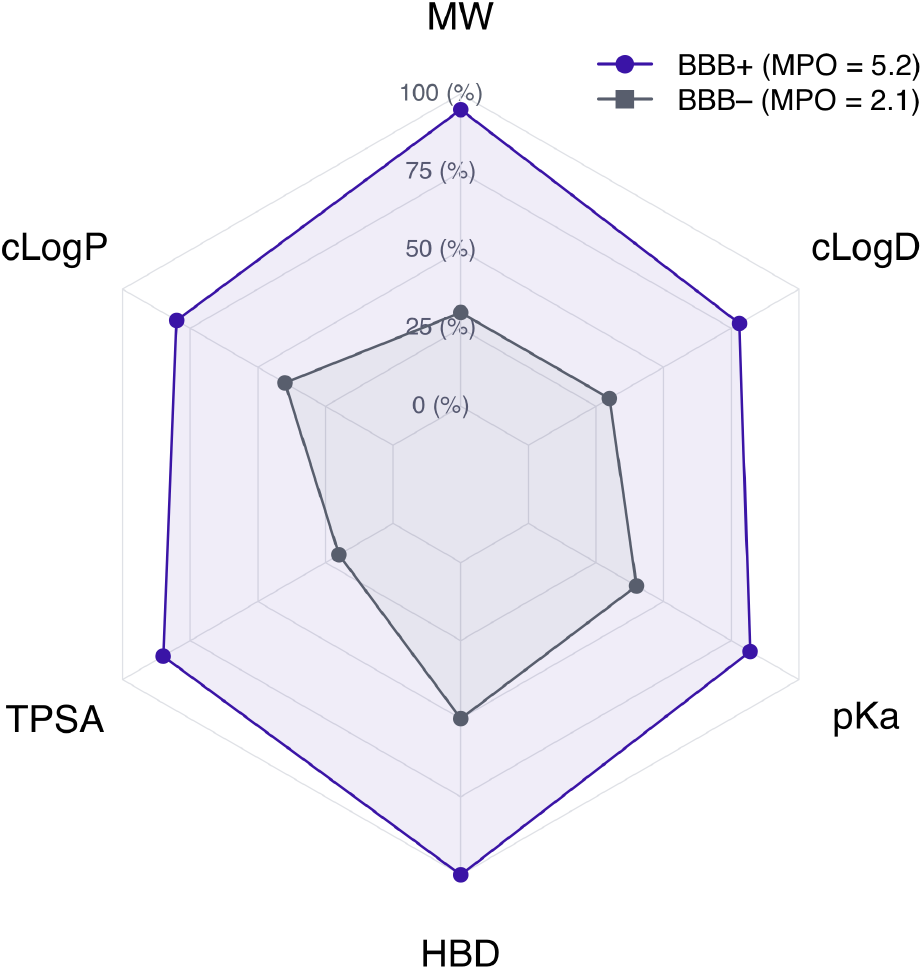
Radar chart comparing mean physicochemical properties of BBB+ (teal) and BBB*−* (purple) compounds across the six CNS-MPO dimensions. BBB+ compounds exhibit lower MW, TPSA, and HBD, and higher cLogP.

### 4.5 Efflux transporter models

Dedicated Random Forest classifiers are trained for each of seven efflux transporters: MDR1 (P-gp/ABCB1), ABCG2 (BCRP), MRP1 (ABCC1), MRP2 (ABCC2), MRP4 (ABCC4), MATE1 (SLC47A1), and OAT3 (SLC22A8). Training labels are derived from ChEMBL bioactivity data, where compounds are labeled as substrates if IC_50_*<* 10 *µ*M or efflux ratio *>* 2.0 in bidirectional transport assays, and as non-substrates otherwise, following established thresholds from the transporter literature.

Each model is trained on a concatenation of 2048-bit ECFP4 (Morgan radius 2) and 166-bit MACCS fingerprints. Models output a substrate probability *P* (substrate) per transporter per compound. If any transporter’s *P* (substrate) exceeds a per-protein threshold calibrated on the ChEMBL validation split, the compound’s final *P*_BBB_is penalized, reflecting the biological principle that a single high-affinity efflux interaction can negate otherwise favorable BBB properties.

On GPUs, a structure-based protein-ligand interaction model provides binding affinity predictions for each compound against the nine efflux transporter protein structures, yielding a second strategy (S3) whose predictions are combined with the fingerprint-based models (S1) into a consensus score. On CPU instances, when the structure-based model is unavailable, the fingerprint-only strategy (S1) provides robust standalone predictions (aggregate AUROC 0.883).

#### 4.5.1 Efflux data pipeline

Training data for the efflux classifiers was curated from ChEMBL (version 33) bioactivity records. For each of the seven target transporters, we extracted compound–target pairs with annotated IC_50_, K_i_, or bidirectional efflux ratio measurements. Compounds were labeled as substrates if IC_50_*<* 10 *µ*M or efflux ratio *>* 2.0, and as non-substrates otherwise. After deduplication by canonical SMILES, the curated dataset comprises 10,803 compound–target pairs across the seven transporters, with per-transporter training set sizes ranging from 1,084 (MATE1) to 2,841 (MDR1).

##### Identifying the barrier proteome

The 65 barrier-functional proteins used in BBB-Nuke were identified through large-scale literature mining rather than manual curation. We collected over 27,000 papers from bioRxiv (19K) and NCBI PubMed (8K) covering BBB permeability, efflux transport, and CNS drug disposition. These papers, comprising over 108 million tokens of scientific text were processed through DeepSeek-based extraction pipelines that converted unstructured findings in prose, tables, supplementary PDFs, and figure captions into structured, machine-readable protein annotations. Extracted protein identifiers were cross-referenced against UniProt to resolve naming ambiguities, map to canonical accessions, and filter to proteins with established expression at the BBB. This process yielded 65 barrier-functional proteins spanning three functional classes: metabolic enzymes (e.g., CYP450 family, MAO-A/B, ACHE), influx/nutrient transporters (e.g., EAAT1–3,LAT1, OATP1A2), and efflux pumps (MDR1, ABCG2, MRP1–5, MATE1, OAT3). UMAP projection of these 65 proteins based on their ChEMBL ligand-binding fingerprint similarity reveals clear functional segregation (Figure 10). Enzymes cluster tightly, reflecting conserved substrate chemotypes within the CYP450 and monoamine oxidase families. Transporters form a more diffuse cluster, consistent with their recognition of structurally diverse solutes across multiple solute carrier families. Efflux proteins cluster separately but proximal to the transporter group, mirroring the biological reality of influx–efflux competition at the barrier. This spatial organization informed the architecture of BBB-Nuke: rather than treating BBB penetration as a single classification problem, the pipeline dedicates one Random Forest classifier per major efflux transporter, treating active exclusion as a first-class predictive signal.

**Figure 10:**
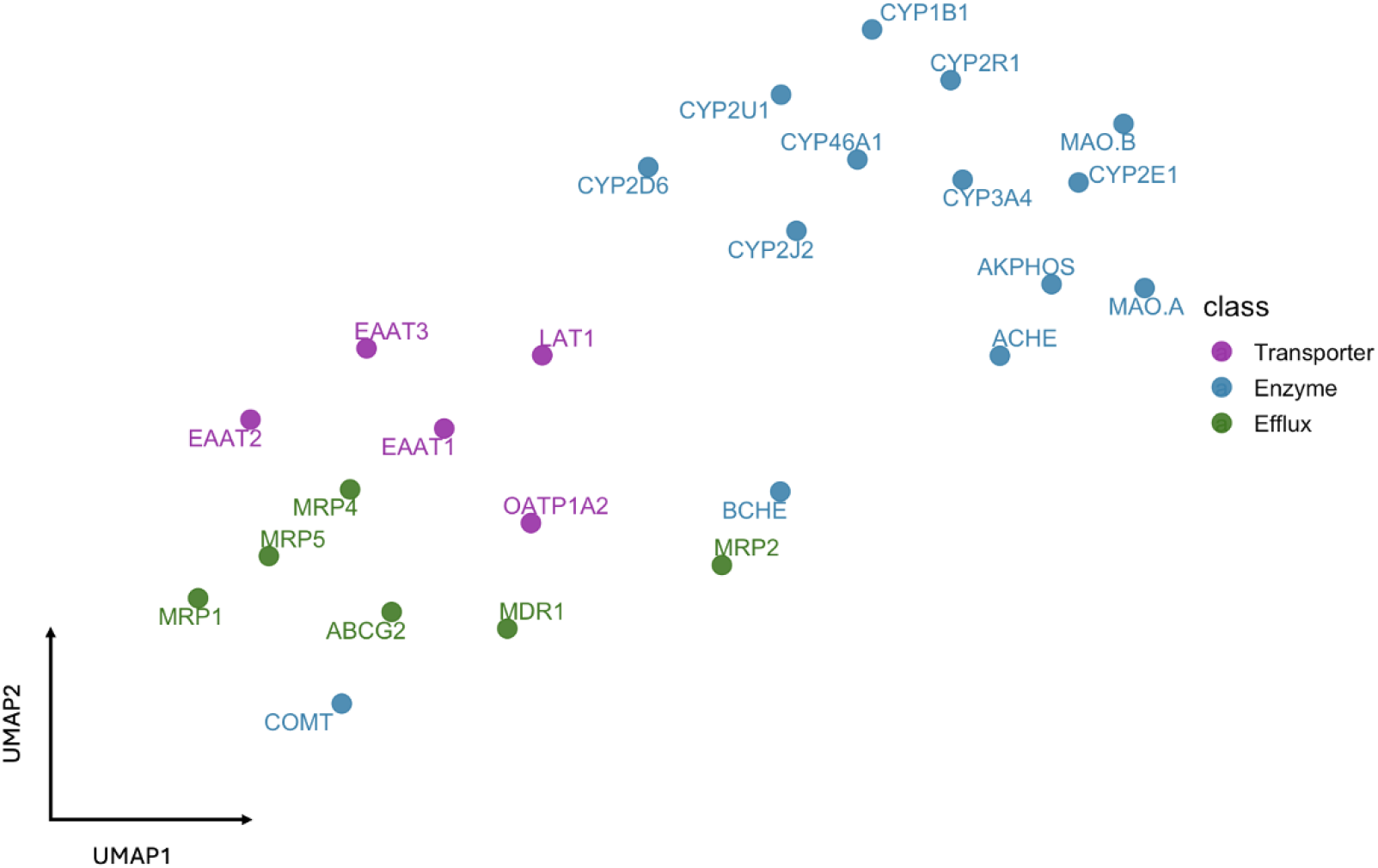
UMAP projection of barrier-functional proteins based on ligand-binding fingerprint similarity. Enzymes (blue), transporters (purple), and efflux proteins (green) exhibit clear functional segregation. Efflux proteins cluster separately and closer to transporters than to enzymes, reflecting the biological relationship between influx and efflux competition at the BBB.

### 4.6 Classifier training

The final BBB classifier is a GradientBoosting model (scikit-learn, 100 estimators, max depth 6, learning rate 0.1). The 67-dimensional input vector is constructed by concatenating the ten physicochemical descriptors, seven efflux substrate probabilities (from the S1 models), and fifty principal components from PCA applied to 2048-bit ECFP4 fingerprints. All features are standardized (zero mean, unit variance) prior to training.

The training set comprises 9,262 compounds from three sources: B3DB [Meng et al., 2021] (7,807 compounds), Benchmark2 (985 compounds), and Benchmark2.5 (470 compounds), deduplicated by canonical SMILES. Labels are binary (BBB+ / BBB*−*). Evaluation uses stratified five-fold cross-validation. The decision threshold is set at 0.30 (*P*_BBB_*≥* 0.30 for BBB+ prediction), calibrated to balance sensitivity and specificity on external benchmarks.

### 4.7 Benchmarking

Three benchmark datasets are used for validation: B3DB [Meng et al., 2021], Benchmark2, and Benchmark2.5. Cross-validation is performed on the combined training set; Benchmark2.5 is additionally evaluated as a fully held-out test set (never included in training, feature engineering, or threshold calibration).

Head-to-head comparisons with CNS-MPO [Wager et al., 2016], LightBBB [Shaker et al., 2021], ADMETlab 2.0 [Xiong et al., 2021], and BBB-Score [Gupta et al., 2019] use published scores or reproduced predictions on the same compound sets. Comparison metrics are AUROC and accuracy at each model’s recommended decision threshold.

### 4.8 Large-scale screening infrastructure

Billion-compound screening was conducted on Azure Machine Learning using 8 A100 GPUs instances and E64ds_v4 instances (64 vCPU, 504 GB RAM). Compounds were processed in chunks of 100,000 with batched GCN inference at mini-batch size 4,096 and vectorized physicochemical computation. Results were written to Apache Parquet format for downstream analysis. Two screening campaigns were executed: the initial campaign processed 365 million compounds from Enamine REAL and PubChem; the second campaign extended coverage to over one billion total compounds.

Post-screening analysis included: Product Quantization k-means clustering (9,971 clusters) on MQN descriptors, UMAP projections of ECFP4 fingerprints (2 million sampled compounds), and TMAP [Probst and Reymond, 2020] tree-map visualizations using MHFP-1024 fingerprints rendered via the Faerun framework [Probst and Reymond, 2018]. All visualizations are generated programmatically via the BBB-Nuke CLI.

## 5 Data Availability

The one-billion-compound screening results are available as an open-access dataset on the Hugging Face Hub `(ATTN-Lab/bbbnuke-screening-1B)`. The dataset contains scored compounds in Apache Parquet format with fields including SMILES, *P*_BBB_, CNS-MPO score, physicochemical properties, and efflux substrate probabilities. The B3DB training dataset is publicly available [Meng et al., 2021]. Efflux transporter ground truth was derived from ChEMBL bioactivity data (version 33).

## 6 Code Availability

BBB-Nuke is available as an open-source Python package. The pipeline can be accessed through three interfaces: a command-line tool `(bbnuke run, bbnuke screen, bbnuke tmap)`, a REST API `(POST /v1/score, GET /v1/proteins, GET /v1/health)`, and a Model Context Protocol (MCP) server for integration with AI assistants. An enterprise desktop edition is available for organizations requiring on-premise deployment with no external network access.

